# γδ T cells do not contribute to peripheral inflammatory pain

**DOI:** 10.1101/390294

**Authors:** Jelena Petrović, Jaqueline Raymondi Silva, Julia P. Segal, Abigail S. Marshall, Cortney M. Haird, Ian Gilron, Nader Ghasemlou

**Affiliations:** Departments of Biomedical & Molecular Sciences, Queen’s University, Kingston, Ontario, Canada; Departments of Anesthesiology & Perioperative Medicine, Queen’s University, Kingston, Ontario, Canada; Departments of Centre for Neuroscience Studies, Queen’s University, Kingston, Ontario, Canada

## Abstract

Circulating immune cells, which are recruited to the site of injury/disease, secrete various inflammatory mediators that are critical to nociception and pain. The role of tissue-resident immune cells, however, remains poorly characterized. One of the first cells to be activated in peripheral tissues following injury are γδ T cells, which serve important roles in infection and disease. Using a transgenic mouse line lacking these cells, we sought to identify their contribution to inflammatory pain. Three distinct models of inflammatory pain were used: intraplantar injection of formalin and incisional wound (as models of acute inflammatory pain) and intraplantar injection of complete Freund’s adjuvant (as a model of chronic inflammatory pain). Our results show that absence of these cells does not alter baseline sensitivity, nor does it result in changes to mechanical or thermal hypersensitivity after tissue injury. These results were consistent in both male and female mice, suggesting that there are no sex differences in these outcomes. This comprehensive characterization suggests that γδ T cells do not contribute to basal sensitivity or the development and maintenance of inflammatory pain.

## Introduction

Celsus assigned four cardinal signs of inflammation in the first century BC: *rubor* (redness), *calor* (heat), *tumor* (swelling), and *dolor* (pain); the Greek physician Galen added *functio laesa* (loss of function) in the 2^nd^ century AD, a critical feature of inflammatory pain that may allow for healing and recovery by promoting post-injury immobility. Pathological inflammatory pain, however, serves no protective purpose; thus, identifying underlying mechanisms of disease will be critical to the development of new therapeutic options. The immune and nervous systems are intimately connected, particularly during inflammatory pain: immune cells and their secreted mediators act on nociceptors in the periphery while neurons, as a consequence, can modulate the inflammatory response [21; 35; 62]. Peripheral inflammation resulting from tissue injury, arthritis, and certain autoimmune diseases is brought on by the well-orchestrated recruitment and activation of circulatory and tissue-resident immune cells, including mast cells, neutrophils and macrophages [34; 66]. These cells and their secreted mediators can alter nociceptor function and activity to induce nociceptor activation and/or peripheral sensitization, triggering an increased responsiveness to noxious stimuli and pain hypersensitivity [21; 34; 35; 62; 66; 68; 85]. Under these conditions of peripheral sensitization, immune cell recruitment and activity can be altered by the neurogenic response from sensory neurons through the release of neuropeptides, neurotransmitters, and cytokines/chemokines [11; 68]. Thus, bi-directional interactions between immune cells and nociceptors are essential in the pathophysiology of inflammatory pain. However, these interactions, as well as the cells and mediators controlling these pain outcomes, remain poorly understood.

Many groups have begun to elucidate these complex neuro-immune interactions. Cytokines (interleukin-1β, tumour necrosis factor), growth factors (nerve growth factor), lipids (prostaglandins), neuropeptides (substance P, calcitonin gene-related peptide) and other inflammatory mediators have been shown to bring about peripheral sensitization and pain hypersensitivity [11; 21; 34; 35; 53; 69]. While the majority of this work has focused on circulating mediators, recent studies have identified specific contributions of immune cell subsets in mediating this pain sensitivity using animal models of injury and disease [5; 6; 19; 43; 44; 52; 70; 80]. These studies have found differing effects for various immune cells. Circulatory cells, including neutrophils and macrophages, have been shown by some groups to modulate inflammatory pain hypersensitivity [5; 6; 19; 33; 70; 80]. The role of tissue-resident cells on the other hand is less well known. Recent work using a transgenic mouse specifically lacking mast cells suggests that these cells have little to no effect on inflammatory pain hypersensitivity [44]. This was a surprising result given that mast cells are known producers of inflammatory mediators including cytokines, growth factors, and inflammatory mediators such as histamine that are known to alter hypersensitivity to stimuli [7; 77; 87]. This brings about the question of whether other tissue-resident immune cells may play a role in inflammatory pain and peripheral sensitization. To understand the role of T cells in inflammatory pain, there are two populations of these cells to consider: circulatory αβ T cells and tissue-resident γδ T cells, which are distinguished by their T cell receptors. In a previous study from our group, we evaluated the depletion of only αβ T cells (using the TCRβ^−/−^ strain) and observed no role for this cell population in inflammatory pain. While αβ T cells have not been shown to modulate inflammatory pain, they do play an important role in neuropathic pain with sexually dimorphic outcomes [47; 48; 74]. γδ T cells are less abundant than αβ T cells, are the primary T cell population found in the gut mucosa and skin, and are absent in TCRδ^−/−^ mice.

Tissue-resident γδ T cells are considered to have qualities of both the innate and adaptive immune systems playing an important role in tissue surveillance, homeostasis and wound repair [27; 31; 60; 79; 83]. γδ T cells respond within minutes to tissue injury, secreting inflammatory mediators such as growth factors and cytokines, are capable of serving as antigen presenting cells, and are critical for host immune defense against infections and autoimmune diseases (e.g., inflammatory bowel disease, multiple sclerosis, and rheumatoid arthritis). Recent work has shown that sensory neurons can suppress γδ T cell numbers [1] and their production of specific cytokines [37], while others have shown γδ T cells to modulate nerve regeneration [42]. Whether γδ T cells can in turn alter nociceptor function/activity remains unknown. We therefore sought out to identify what role γδ T cells might play in baseline sensitivity, due to their close proximity to sensory fibres in the skin, and the response to inflammatory pain using TCRδ^−/−^ mice, which lack these cells. Acute and persistent peripheral inflammation were used to mimic human clinical inflammation: the intraplantar injection of formalin (non-reflexive, spontaneous pain) or complete Freund’s adjuvant (chronic inflammatory pain), and plantar incisional wound (acute post-surgical pain) [23; 59].

## Materials and Methods

All protocols were approved by the Queen’s University Animal Care Committee and followed the ARRIVE guidelines [40] and those of the Canadian Council on Animal Care. All surgical and behavioural work was carried out between 9am to 5pm in a facility where lights are on between 7am to 7pm.

### Animals

B6.129P2-Tcrd^tm1Mom^/J mice (TCRδ^−/−^; Jackson Laboratory, Bar Harbor, ME), which lack γδ T cells [30], were backcrossed to C57BL/6J mice (Jackson Laboratory) to generate TCRδ^+/−^ F1 progeny. Heterozygous mice from different parents were subsequently used to establish the colony including wildtype, heterozygous and knockout littermates. All experiments were carried out using adult mice between 6-12 weeks of age, housed at a maximum of four per cage on a 12-hour light/dark cycle in a temperature- (21°C (±1°C)) and humidity-controlled room, with food and water provided *ad libitum.* The colony was maintained and genotyped by an independent experimenter, ensuring that all work was carried out blinded to genotype. Wildtype, heterozygous, and knockout littermates were used in all experiments, with at least two independent replications.

### Formalin

Male (n=6-9) and female (n=8-13) TCRδ littermates received intraplantar injections with 20μl of a 5% formalin solution [diluted from a 37% formaldehyde stock solution (Sigma-Aldrich, St. Louis, MO) re-suspended in 0.9% NaCl solution (Hospira, Saint-Laurent, QC)] using a Hamilton syringe with 27-gauge needle, as previously described [78].

### Plantar incisional wound

Male (n=7-10) and female (n=8-12) TCRδ littermates were anesthetized with 2% (v/v) isoflurane USP (Fresenius Kabi Canada, Toronto, ON) and the left hindpaw sterilized three times sequentially with 10% (w/v) Proviodine^®^ Povidone Iodine Solution (Rougier Pharma, Mirabel, QC) and 70% (v/v) ethanol. A 5-8mm incision was made using a #11 scalpel from the base of the heel to the first walking pad, cutting through the skin and underlying muscle along the midline of the plantar surface, as previously described [19]. The wound was closed at two sites with 6-O silk sutures (Ethicon, Cincinnati, OH). Pain hypersensitivity was measured post-incision at 3 and 6 hours, and 1, 2, 3, 4 and 7 days.

### Complete Freund’s adjuvant

Male (n=7-10) and female (n=8-12) TCRδ littermates received intraplantar injections with 20μl of complete Freund’s adjuvant (Sigma-Aldrich) using a Hamilton syringe with 27-gauge needle, as previously described [19]. Mice were not anesthetized for the injections and were immediately returned to their home-cage until behavioural experiments were carried out. Pain hypersensitivity was measured at 3 and 6 hours, and 1, 2, 3, 4 and 7 days following plantar CFA injection.

### Behavioural assays

Mice injected with formalin were immediately placed singly in clear glass chambers and nociceptive behaviour (characterized by licking and biting of the injected hindpaw) was observed over 60 minutes and recorded at 5-minute intervals. Behavioural analysis was further divided into two phases: an acute phase lasting 0-10 minutes and a tonic phase from 10-60 minutes. Mice undergoing intraplantar injection of CFA or plantar incisional wound were habituated to the equipment and experimenter over five consecutive days prior to all behavioural testing with at least 30 minutes in each experimental apparatus. Baselines measurements were then taken on three separate days and the average response calculated. All behavioural assays were carried out using at least two independent cohorts of mice by a researcher blinded to animal genotype using methods previously described by our group [19] and briefly outlined below.

#### von Frey mechanical sensitivity assay

Mice were placed in polycarbonate boxes on a wire mesh with black dividers to reduce interactions between animals. The left hindpaw was stimulated with graded von Frey monofilaments (North Coast Medical, Gilroy, CA). Gentle stimulation with these monofilaments were used to measure the lowest force (in grams) at which the mouse responds at least 50% of the time (paw threshold response), as previously described [19].

#### Acetone cold sensitivity test

Similar to the von Frey assay, the acetone test is performed by placing mice on the same wire mesh apparatus. A small drop of acetone was gently applied to the left hindpaw using a 1cc syringe (Becton Dickinson, Franklin Lakes, NJ) and the amount of time (in seconds) spent licking and biting the affected paw was measured. At least two measurements were recorded and averaged per timepoint, with a minimum of five minutes between each stimulation.

#### Hargreaves radiant heat sensitivity test

Mice were assessed for thermal heat hypersensitivity by performing the Hargreaves radiant heat test [86]. Mice were placed in individual transparent polycarbonate compartments on a heated glass base set to a constant temperature of 30°C (±1°C) (IITC Life Science, Woodland Hills, CA). A radiant heat source was focused onto the plantar surface of the left hindpaw, creating a 4×6 mm radiant heat source for stimulation, and the latency to withdrawal measured. A maximum cut-off of 30 seconds was used to prevent tissue damage. Three measurements were averaged from each timepoint.

#### Hot/cold plate test

Thermal sensitivity was assessed at specific temperatures using an air-cooled thermoelectric plate (TECA Corporation, Chicago, IL). The plate was set to 0, 50, 52, or 55°C, allowed to stabilize at the temperature for 15 minutes, and animals placed in the center of the platform. The time to first response (e.g., fast paw withdrawal, shaking, licking/biting) and jump (all four paws lifted off the plate) was recorded. A maximum cut-off of 30 seconds was used to prevent tissue damage. Only one measurement was taken at each temperature per day to prevent learning behaviours; mice that exhibited learning behaviours (e.g., scaling the enclosure) were excluded from analysis [44].

### Immunohistochemistry

Immunostaining for γδ T cells was carried out as described elsewhere (Marshall, Silva, Gilron, and Ghasemlou, in preparation). Briefly, mice were deeply anesthetized and sacrificed by transcardial perfusion with 2% paraformaldehyde in 0.1M phosphate buffer. Ears were removed, post-fixed for 1 hour, and cryoprotected in 30% sucrose. Serial cryostat sections (15μm thickness) were obtained for histological analysis. Samples were incubated with hamster anti-mouse TCRδ (1:100; Invitrogen, Waltham, MA), followed by goat anti-hamster IgG conjugated to FITC (1:200; BioLegend, San Diego, CA). Slides were coverslipped using Vectashield mounting medium with DAPI (Vector Labs, Burlingame, CA) and visualized using an AxioSkop2 fluorescent microscope and AxioVision software (Carl Zeiss AG, Jena, Germany).

### Statistical analysis

All statistical analyses were carried out using SigmaPlot version 11.0 software package (Systat Software, San Jose, CA). Data are expressed as mean ± standard error of the mean (SEM) throughout the text and in figures. One-way analysis of variance (ANOVA) was used for a direct comparison between two or more groups, while a two-way repeated-measures ANOVA was used to assess the effect of time between two or more groups. *Post-hoc* Tukey tests were used where appropriate with significance set at P<0.05.

## Results

### γδ T cells do not contribute to baseline thermal or mechanical sensitivity

Immunohistochemical analysis was used to visualize presence/absence of these cells in TCRδ^+/+^ and TCRδ^−/−^ mice to ensure complete absence of γδ T cells in null mice. As expected, γδ T cells were present in the epidermal layer of TCRδ^+/+^ mice, yet absent in TCRδ^−/−^ littermates (Figure 1A). We began our work by assessing whether loss of γδ T cells causes a change to either baseline mechanical or thermal sensitivity in male and female mice, as these cells are resident in the skin and other barrier organs and could interact with sensory fibres during development. Mechanical sensitivity, measured as the 50% threshold using graded von Frey monofilaments, did not show any difference between TCRδ^+/+^, TCRδ^+/−^, and TCRδ^−/−^ mice in either males (Figure 1B; n=17-22 per genotype; one-way ANOVA, P=0.402) or females (Figure 1B; n=15-18 per genotype; one-way ANOVA, P=0.276). Mechanical threshold was slightly, but not significantly, increased in TCRδ^−/−^ males relative to wild-type (1.39±0.12g vs. 1.21±0.09g, respectively), though female mice did not exhibit such an effect.

**Figure 1.**
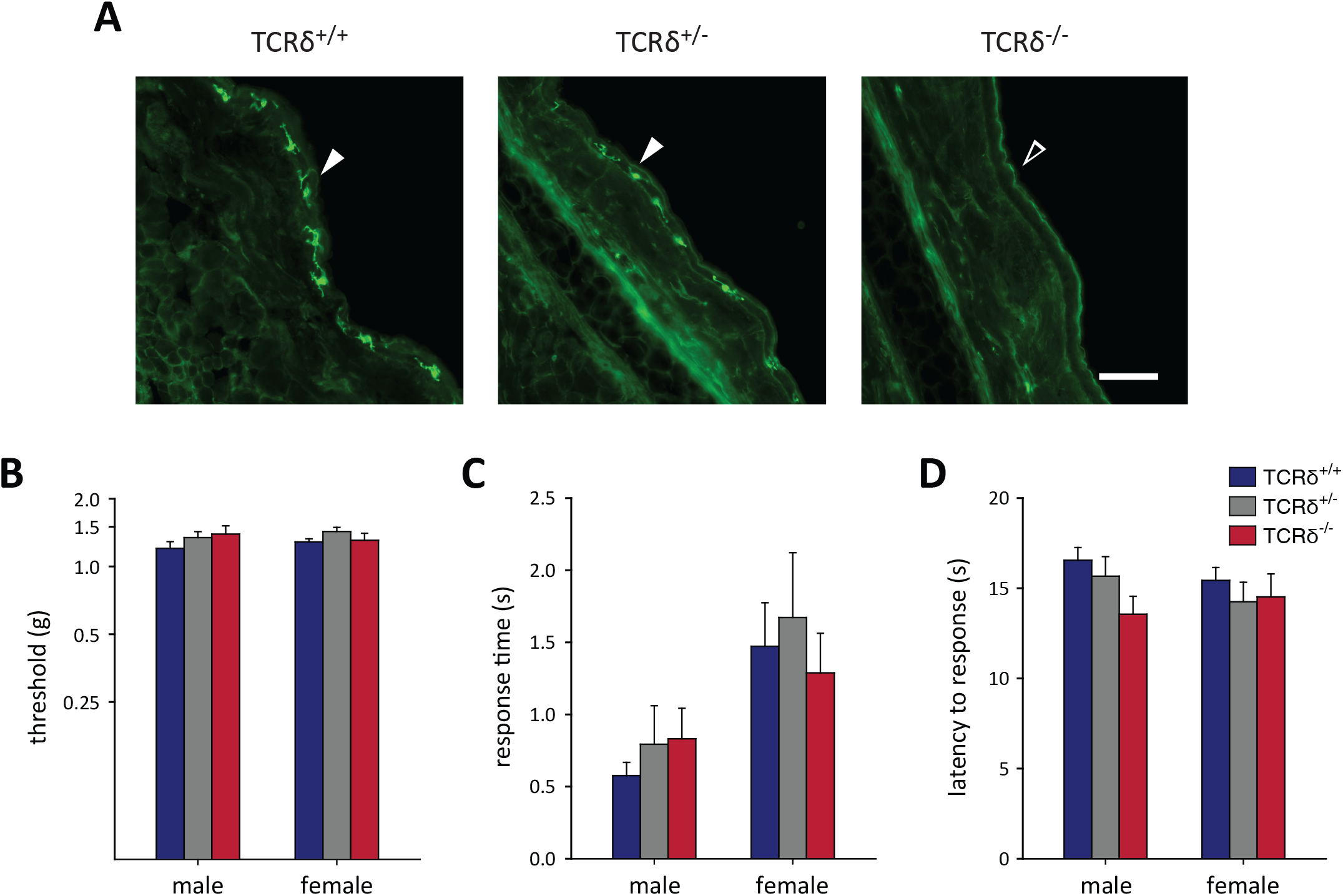
Absence of γδ T cells does not affect basal mechanical or thermal sensitivity. (**A**) Sections of the ear from wildtype, heterozygous, and knockout TCRδ mice immunostained for γδ T cells using an antibody recognizing the δ T cell receptor subunit. Representative micrographs show γδ T cells are present in TCRδ^+/+^ and TCRδ^+/−^ mice, but not in TCRδ^−/−^ littermates. (**B**) Mechanical thresholds, measured as the von Frey monofilament corresponding to a 50% response, is not affected by loss of γδ T cells in male (P=0.402, one-way ANOVA; n=17-22 per genotype) or female (P=0.276, one-way ANOVA; n=15-18 per genotype) mice. (C) Cold thermal responses were assessed using the acetone test, measured as total response time (e.g., licking and biting of the affected hindpaw), was not different between male (P=0.669, one-way ANOVA; n=14-16 per genotype) or female (P=0.758, one-way ANOVA; n=10-15 per genotype) littermates. (D) Thermal heat hypersensitivity was measured as the latency to response following stimulation of the hindpaw by a radiant heat source. No differences were observed in either male (P=0.086, one-way ANOVA; n=17-19 per genotype) or female (P=0.679, one-way ANOVA; n=15-18 per genotype) mice. Graphs show mean ± SEM, scale bar=50μm.

We next measured responses to thermal stimuli at baseline. Cold sensitivity was assessed using the acetone test and heat sensitivity was measured using the Hargreaves radiant heat test. The response time to application of acetone to the hindpaw did not show a significant effect in either males (Figure 1C; n=14-16 per genotype; one-way ANOVA, P=0.669) or females (Figure 1C; n=10-15 per genotype; one-way ANOVA, P=0.758). The response to the radiant heat source was not different between the three genotypes in either male (Figure 1D; n=17-19 per genotype; one-way ANOVA, P=0.086) or female mice (Figure 1D; n=15-18 per genotype; one-way ANOVA, P=0.679). Although the response time was slightly faster in male TCRδ^−/−^ mice relative to wild-type littermates (13.55±1.00s vs. 16.55±0.71s, respectively), such an effect was not evident in female mice. The cold and hot plate test was used to identify whether there were any differences in noxious thermal response between the three genotypes that could not be measured using the Hargreaves and acetone tests, using fixed temperatures between 0 to 55°C. As before, male TCRδ^+/+^ (n=7-11), TCRδ^+/−^ (n=8-15), and TCRδ^−/−^ (n=6-12) mice did not show any significant differences at 0, 50, 52, or 55°C in either time to first response (e.g. flinching of the hindpaw; Figure 2A; one-way ANOVA, P≥0.193) or first jump (Supplemental Figure 1A; oneway ANOVA, P≥0.444). Female TCRδ^+/+^ (n=12-17), TCRδ^+/−^ (n=6-13), and TCRδ^−/−^ (n=6-13) mice also did not show any differences in response time to flinch at 0, 50, or 52°C (Figures 2B; one-way ANOVA, P≥0.099). While there was a group effect between the three strains at 55°C for latency to first response (one-way ANOVA, P=0.039),*post-hoc* analysis did not show significant differences between the female strains at 55°C (e.g., TCRδ^+/+^ vs TCRδ^−/−^, TCRδ^+/−^ vs TCRδ^−/−^, etc.; one-way ANOVA, with *post-hoc* Tukey test P=0.063). Time to first jump was not significant at any temperature in female mice (Supplemental Figure 1B; one-way ANOVA, P≥0.129). Thus, loss of γδ T cells does not appear to alter basal mechanical or thermal sensitivity.

**Figure 2.**
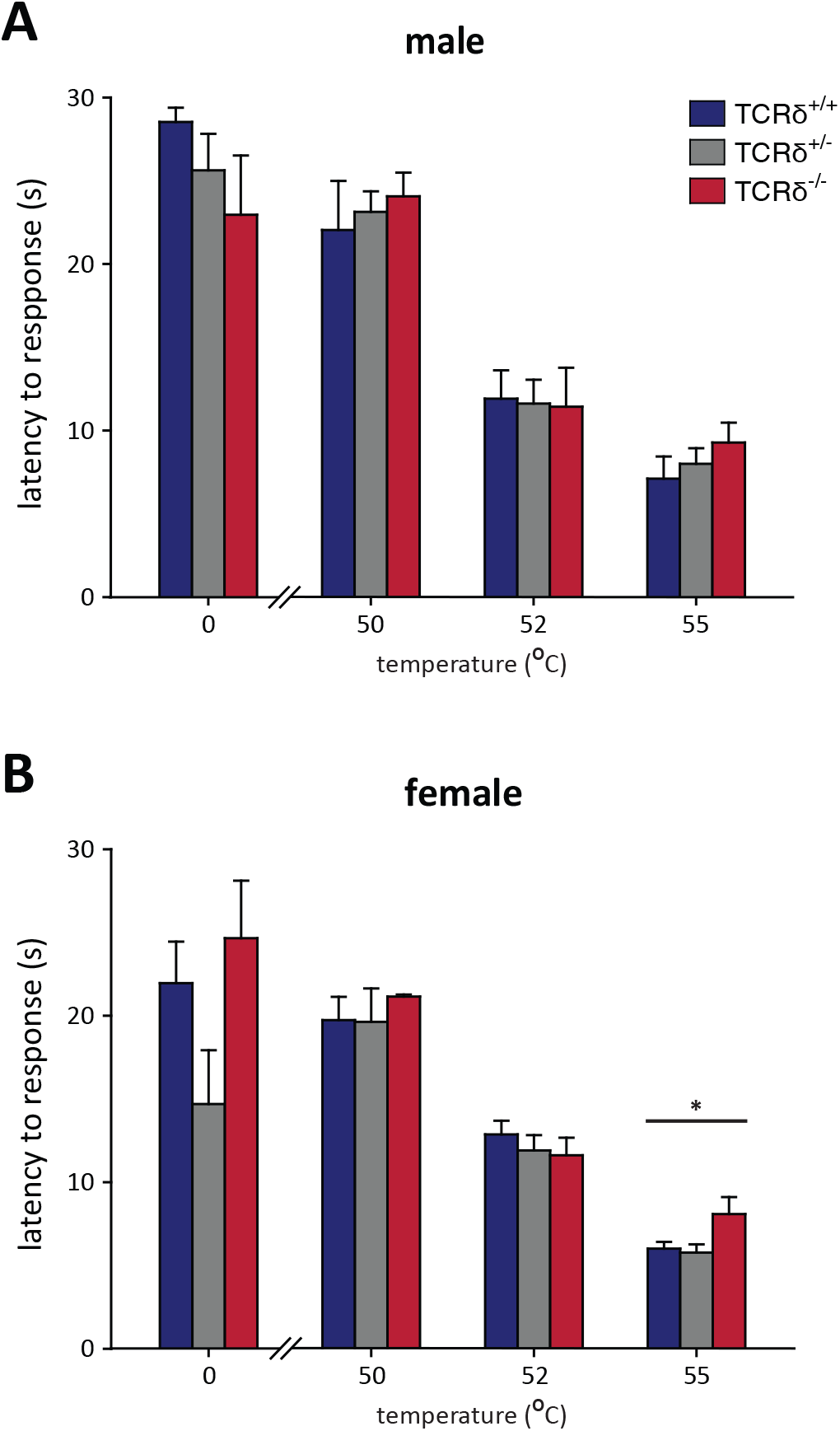
Response to noxious heat and cold are unaffected by loss of γδ T cells. (**A**) No differences were observed in latency to paw withdrawal (e.g., flinch) using the hot and cold plate test in male TCRδ littermates (P P≥0.193, one-way ANOVA; n=6-15 per genotype) at any of the temperatures examined. (**B**) Female mice assessed for latency to first response did not exhibit differences at 0, 50, or 52°C (P≥0.099, one-way ANOVA; n=6-17 per genotype). While there was a significant group effect for genotype at 55°C (*P=0.039, one-way ANOVA), *post-hoc* Tukey analysis was not significant between the three groups (P≥0.063).

### γδ T cells do not contribute to inflammatory pain

Specific circulating and skin-resident immune cells have been found to control inflammatory pain responses. The contribution of γδ T cells to the inflammatory pain response was assessed using standard assays, where all experiments were conducted using at least 2 cohorts of littermates.

#### Formalin

We first assessed the contribution of γδ T cells to acute inflammatory pain outcomes using the formalin test [72; 78], where nociceptive pain, measured as time spent licking/biting the affected hindpaw, often lasts for approximately 60 minutes. Male TCRδ mice (n=6-9 per genotype) did not show an effect across the three genotypes studied over the course of their response (Figure 3A; two-way repeated measures ANOVA, P=0.403). Differences were also not observed when the response times were divided into acute (0-10min) and tonic (10-60min) phases (Figure 3B; one-way ANOVA, p≥0.400). Female mice (n=8-13 per genotype) similarly do not show an effect over the duration of their response (Figure 3C; two-way RM-ANOVA, P=0.353), or over the acute or tonic phases (Figure 3D; one-way ANOVA, p≥0.338). This suggests that γδ T cells do not contribute to formalin-induced inflammatory pain.

**Figure 3.**
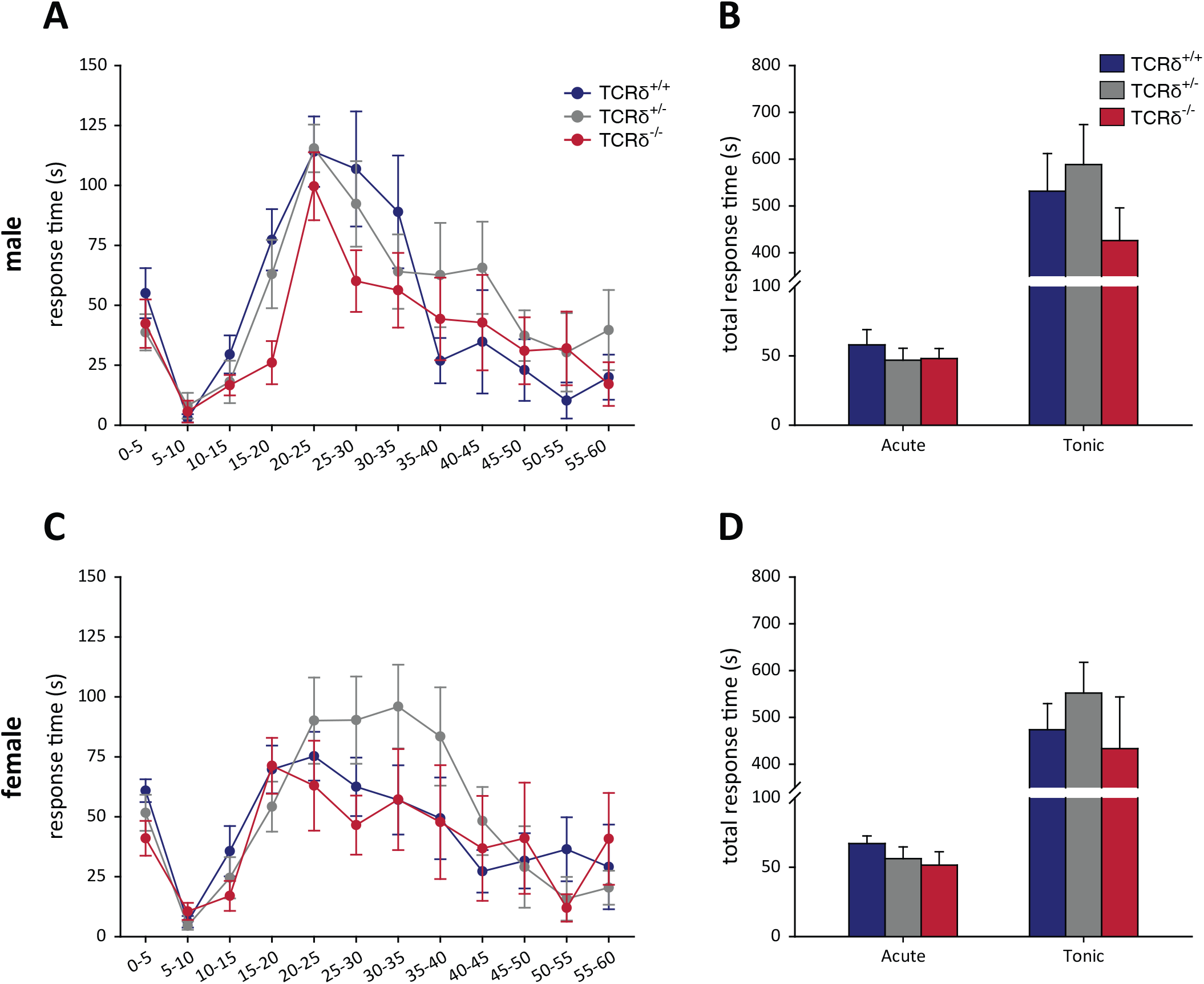
Response to formalin is unaffected by absence of γδ T cells. TCRδ littermates were injected with formalin and the response time measured over 60 minutes. Male mice (n=6-9 per genotype) did not show an effect over the duration of response (**A**; P=0.403, two-way RM-ANOVA) or during acute and tonic phases (**B**; P≥0.400, one-way ANOVA). Female mice (n=8-13 per genotype) also did not show a significant effect over the duration of response (**C**; P=0.353, two-way RM-ANOVA) or in acute/tonic phases (**D**; P>0.338, one-way ANOVA).

#### Incisional wound

We next considered the contribution of γδ T cells to the development and maintenance of acute inflammatory pain (lasting 2-4 days after injury) following plantar incisional wound, a model of post-surgical pain. Mechanical hypersensitivity in the injured hindpaw showed no difference between male TCRδ^+/+^, TCRδ^+/−^ and TCRδ^−/−^ mice (n=7-10 per genotype, P=0.064, two-way RM-ANOVA; Figure 4A). Similarly, hypersensitivity to heat stimuli also showed no significant differences (P=0.215, two-way RM-ANOVA, Figure 4B). Female TCRδ mice (n=8-12 per genotype) also did not show any significant effects after incisional wound for either mechanical hypersensitivity (P=0.942, two-way RM-ANOVA, Figure 4C) or thermal hypersensitivity (P=0.675, two-way RM-ANOVA, Figure 4D), suggesting γδ T cells do not affect post-surgical inflammatory pain.

**Figure 4.**
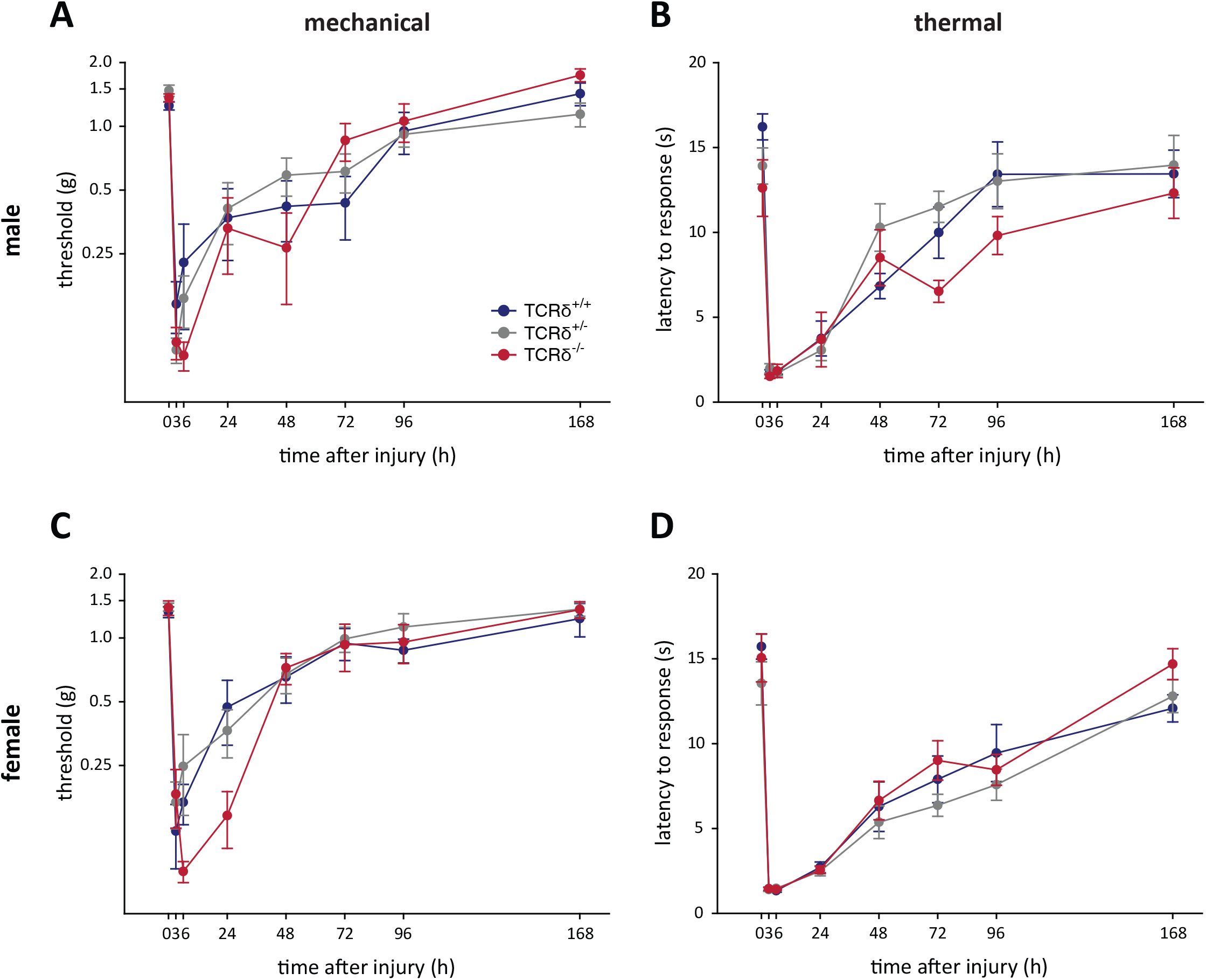
γδ T cells do not contribute to mechanical and thermal hypersensitivity after incisional wound. Male TCRδ littermates (n=7-10 per genotype) did not exhibit differences in mechanical thresholds (**A**; P=0.064, two-way RM-ANOVA), measured with von Frey monofilaments, or heat hypersensitivity (**B**; P=0.215, two-way RM-ANOVA), measured as the latency of response to a radiant heat stimulus. A similar effect was observed in female TCRδ littermates (n=8-12 per genotype) for both mechanical (**C**; P=0.942, two-way RM-ANOVA) and thermal (**D**; P=0.675, two-way RM-ANOVA) hypersensitivity.

#### Complete Freund’s adjuvant

To determine whether γδ T cells contribute to chronic inflammatory pain, we carried out intraplantar injection of CFA. Mechanical and thermal hypersensitivity do not return to baseline levels following intraplantar CFA injection as occurs following incisional wound, providing an opportunity to assess the contribution of these cells to a prolonged inflammatory response. Similar to the formalin and incisional wound models, no significant effects were observed in male (n=9-12 per genotype) or female (n=6-9 per genotype) TCRδ littermates when assessed for mechanical (P=0.226 [male], P=0.530 [female], two-way RM-ANOVA; Figures 5A, C) and thermal hypersensitivity (P=0.943 [male], P=0.857 [female], two-way RM-ANOVA; Figures 5B, D). Thus, γδ T cells likely do not contribute to chronic inflammatory pain outcomes.

**Figure 5.**
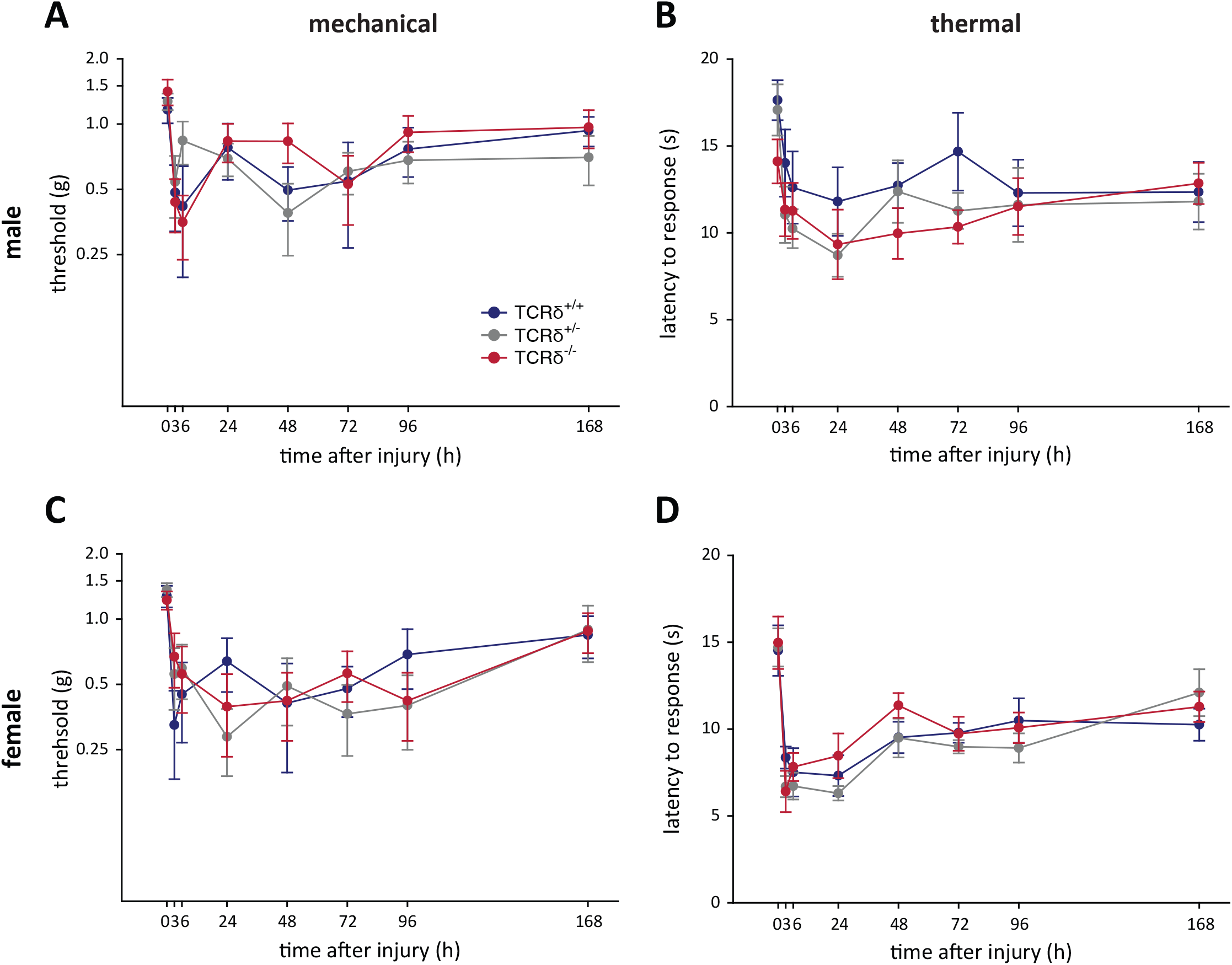
CFA-induced hypersensitivity is unaffected by loss of γδ T cells. TCRδ littermates received intraplantar injections of complete Freund’s adjuvant and pain outcomes were measured over 7 days. Differences in mechanical (**A**; P=0.226, two-way RM-ANOVA) or thermal (**B**; P=0.943, two-way RM-ANOVA) in male mice (n=9-12 per genotype). Female littermates (n=6-9 per genotype) also did not show any differences in mechanical (**C**; P=0.530, two-way RM-ANOVA) or thermal (**D**; P=0.857, two-way RM-ANOVA) responses.

## Discussion

Inflammatory pain is associated with tissue damage, infection, or autoimmune disease, as well as the presence of immune cells that can contribute to pain outcomes and wound healing. This pain often serves an important role in allowing healing and recovery to occur by promoting temporary immobility, such as after injury or infection. However, it can also be pathological in nature as it transitions into persistent pain where it serves no purpose, as is often the case in autoimmune diseases and in many patients following surgery once their wounds have healed [28; 38; 39]. Understanding how the immune cells and their mediators contribute to the development and maintenance of pain will be crucial to the development of safe and efficacious therapeutics for the treatment of inflammatory pain.

We therefore set out to determine the contribution of γδ T cells to the development and maintenance of acute and chronic inflammatory pain, beginning by first backcrossing the B6.129P2-TCRd^tm1Mom/J^ mice to the C57BL/6J background, in an effort to account for genetic background-dependent variations in behavioural phenotypes [41; 57; 58]. We began by backcrossing B6.129P2-T*crd*^*tmlMom*^/J mice with wildtype C57BL/6J mice to generate TCRδ^+/+^, TCRδ^+/−^, and TCRδ^−/−^ littermates that were used in all studies, resulting in a total of 14 backcrosses for this strain. All experiments were carried out using both male and female mice in an effort to identify any effects controlled by sex. We first found that loss of these cells did not have an effect on basal sensitivity, as these cells may interact with sensory fibres in the skin during development and thus affect their function. Using three models of peripheral inflammatory pain, including intraplantar injection of formalin or CFA, and incisional wound, our work revealed that γδ T cells do not alter mechanical or thermal sensitivity. It is important to note that the pain outcomes observed in TCRδ wildtype and heterozygous animals in the formalin, incisional wound, and CFA models matches that observed in C57BL/6 mice in previous studies by our group and others [14; 19; 58]. We previously identified the contribution of myeloid and lymphoid cells to inflammatory pain, using cell-specific strategies to deplete neutrophils, non-neutrophil myeloid, and αβ T cells [19]. While only non-neutrophil myeloid cells were found to alter behavioural outcomes, the role of most tissue-resident cells to inflammatory pain remained unknown.

γδ T cells are a population of skin-resident cells that lie at the intersection of the innate and adaptive immune response and are among the first cells to be activated in the skin following tissue injury or viral/bacterial infection [12; 17; 31; 63; 65; 71]. These cells are known for their dendritic-like projections that allow them to intimately interact with their local environment and neighbouring cells, such as keratinocytes, Langerhans cells, and melanocytes [24; 45]. γδ T cells are similar in appearance to microglia: stationary under homeostatic conditions, with dendritic projections that allow them survey their immediate milieu for signs of injury or infection [13; 22]. While our results suggest that γδ T cells do not contribute to inflammatory pain hypersensitivity, only three models of disease were used. Assessing the role of these cells may show an important effect in pain outcomes following bacterial infection, where they are known to have an effect in both skin [15; 46] and lung [1; 8; 55], or in models of inflammatory bowel disease [16; 29; 36]. We hypothesize this to be possible due to the high number of these cells in the lungs and lining the gut mucosa. The site of injury/infection may be critical, given that γδ T cells are more prevalent in the ear and back skin than in the hindpaw, where their number and morphology are more similar to that observed in human skin [unpublished observations, 4].

γδ T cells participate in the early stages of an immune response by recognizing lipid-ligand antigens, unprocessed proteins, and phosphor-antigens without MHC-restricted presentation [3; 10; 25; 26]. Several studies showed that γδ T cells contributes to the development of an inflammatory response in various tissues (e.g., gut, lungs, spinal cord), through expression of inflammatory cytokines and regulatory factors including interferon (IFN)-γ, IL-17, TNF-α, granzymes, and insulin-like growth factor 1 [10] This can result in the recruitment of circulating immune cells to the site of inflammation [18] and can modulate γδ T cell interaction with other immune cells, including B and αβ T cells [50; 64; 67; 75; 76]. Two major families of mediators secreted by γδ T cells include keratinocyte and fibroblast growth factors (KGFs and FGFs). While several FGF family members have been found to directly activate sensory neurons [49; 62; 73], KGFs are not known to affect sensory neuron activity, though keratinocytes themselves have recently been implicated in modulating nociception [2; 61], as well as itch [51; 81] and mechanosensitivity [9; 56]. Although our results do not show an effect for γδ T cells in the nociceptive response following peripheral inflammation, these cells may still play an important role in itch and other skin pathologies and could prove useful in identifying novel underlying cellular and molecular mechanisms. When evaluating the influence of these cells in the skin, there are some conflicting results in literature. Some studies suggest that γδ T cells are important for the development of the inflammatory response, with mechanisms similar as described above. Other studies have shown that these cells are not required or negatively regulate skin inflammation [20; 84]. These studies have almost exclusively studied γδ T cells in the ear and back, and no studies have been carried out in the footpad.

Besides their potential role in skin inflammation, it is well known that γδ T cells play an important role in wound healing. Skin-resident γδ T cells are called dendritic epidermal T cells (DETCs) when strictly limited in their distribution to the epidermis. Their distribution in the skin epithelium allows the intimate contact with other resident cell types, as well as with the extracellular matrix. They recognize an unidentified antigen expressed by damaged, stressed, or transformed keratinocytes in the epidermis, which results in the production of inflammatory mediators that have specific effects on other epithelial cells [82]. Among these regulatory factors, it has been demonstrated that the insulin-like growth factor-1, KGFs, and FGFs, are involved in tissue repair after injury. In fact, rapamycin treatment of mice, which disables γδ T cell proliferation, migration, and expression of normal levels of growth factors, compromises the wound repair in the injured skin. However, treatment with insulin-like growth factor-1, produced by DETCs, results in restoration of normal wound closure in those animals [54]. Similarly, the addition of DETCs or recombinant KGF in skin organ culture from γδ T cell-deficient mice restored normal wound healing in this tissue [31]. Finally, upon TCR stimulation in vitro,

DETCs produce FGF, which has been suggested to promote keratinocyte proliferation and accelerates wound repair [32]. Altogether, these findings suggest that DETCs and γδ T cells might play an important role in the biological function of wound repair. It has also been suggested that γδ T cells might have regulatory roles in the inflammatory response initiated after tissue injury. For instance, these cells are responsible for the modulation of myeloid cell infiltration after burn-induced wound by suppressing the inflammatory response, leading to the initiation of the proliferative phase of wound healing [65].

While our results demonstrate that γδ T cells do not contribute to inflammatory nociception, this is limited by our use of three specific models – the intraplantar injection of formalin or complete Freund’s adjuvant and incisional wound. While this is the first study of the function of these cells in mediating pain outcomes, our work is limited to inflammation in the hindpaw. We speculate that future work examining the function of γδ T cells in other models of pain/nociception may yet identify a role for these cells.

## Acknowledgements

This work was supported by grants from the Canadian Institutes of Health Research, the J. P. Bickell Foundation, and Queen’s University to NG. The authors thank Dr. Michael D. Kawaja for technical support.

